# Frontal language areas do not emerge in the absence of temporal language areas: A case study of an individual born without a left temporal lobe

**DOI:** 10.1101/2021.05.28.446230

**Authors:** Greta Tuckute, Alexander Paunov, Hope Kean, Hannah Small, Zachary Mineroff, Idan Blank, Evelina Fedorenko

**Author notes:** Corresponding Authors Greta Tuckute or Ev Fedorenko. Conflict of interest The authors declare no competing financial interests.

## Abstract

Language relies on a left-lateralized fronto-temporal brain network. How this network emerges ontogenetically remains debated. We asked whether frontal language areas emerge in the absence of temporal language areas through a ‘deep-data’ investigation of an individual (EG) born without her left temporal lobe. Using fMRI methods that have been validated to elicit reliable individual-level responses, we find that—as expected for early left hemisphere damage—EG has a fully functional language network in her right hemisphere (comparable to that in n=145 controls) and performs normally on language assessments. However, we detect no response to language in EG’s left frontal lobe (replicated across two sessions, 3 years apart). Another network—the multiple demand network—is robustly present in frontal lobes bilaterally, suggesting that EG’s left frontal cortex can support non-linguistic cognition. The existence of temporal language areas therefore appears to be a prerequisite for the emergence of the frontal language areas.

## Introduction

Any typically developing child acquires a language, or multiple languages, in the presence of linguistic input. In the adult human brain, language processing recruits a fronto-temporal network (e.g., Fedorenko & Thompson-Schill, 2014). In most individuals, this network is dominant in the left hemisphere (LH), as evidenced by both i) more robust LH activity in response to language processing as measured with fMRI and other brain imaging techniques (e.g., Petersen et al., 1988; Binder et al., 1997; Vigneau et al., 2006; Fedorenko et al., 2010; Mahowald & Fedorenko, 2016), and ii) a greater likelihood of linguistic deficits that result from the stimulation of (e.g., Penfield & Roberts, 1959; Ojemann et al., 1989) or damage to / degeneration of (e.g., Geschwind, 1971; Damasio, 1992; Benson and Ardila, 1996; Bates et al., 2003; Mesulam et al, 2014; Fridriksson et al., 2016) the LH in adulthood.

How the language network emerges and develops remains poorly understood. An important contributing factor is the difficulty of probing the functional organization of children’s brains between 1 and 3 years of age, when language makes the biggest developmental leap (e.g., Brown, 1973; Frank et al., 2020). A number of studies have examined responses to speech in infant brains (e.g., Dehaene-Lambertz et al., 2002; Pena et al., 2003; May et al., 2011; Perani et al., 2011; Naoi et al., 2013; see Dehaene-Lambertz & Spelke, 2015 for a review), and a now substantial, and growing, literature has examined language brain function in children aged 4-5 and through adolescence (e.g., Balsamo et al., 2002; Ahmad et al., 2003; Blumenfeld et al., 2006; Chou et al., 2006; Wood et al., 2004; Friederici et al., 2011; Nuñez et al., 2011; Langeslag et al., 2013; Xiao et al., 2016; Olulade et al., 2020; see Rosselli et al., 2014 for a review). However, by age 4-5 years, the language network in the dominant hemisphere appears to be largely similar to that of adults (e.g., Wood et al., 2004; Friederici et al., 2011; Berl et al., 2014), although activations tend to be more bilateral at younger ages (e.g., Holland et al., 2001; Chou et al., 2006; Szaflarski et al., 2006; Brauer et al., 2007; Holland et al., 2007; McNealy et al., 2011; May et al., 2011; Berl et al., 2014; Olulade et al., 2020; see Holland et al., 2007 for a review). So the functional architecture of language during the critical window of development remains largely underexplored.

Numerous studies have examined the *anatomy* of cortical areas and white-matter pathways that are plausibly important for language function (e.g., Sowell et al., 2002; Hagmann et al., 2010; Perani et al., 2011; Brauer et al., 2013; Can et al., 2013; Broce et al., 2015, Tak et al., 2016), including during the first few years of life (e.g., Amunts et al., 2003; Pujol et al., 2006; Su et al., 2008; Perani et al., 2011; Tak et al., 2016). However, evidence from studies of anatomy alone is challenging to interpret given high inter-individual variability in the locations of functional language areas (e.g., Steinmetz & Seitz, 1991; Demonet et al., 1993; Ojemann, 1979; Fedorenko et al., 2010; Mahowald & Fedorenko, 2016; Braga et al., 2020), and poor correspondence between those areas and macro-anatomic landmarks (e.g., Fischl et al., 2008; Frost & Goebel, 2011; Tahmasebi et al., 2012).

Some constraints on theories of neural language development have been derived from studies of early brain damage and subsequent reorganization. For example, evidence from early LH damage has established that the right hemisphere (RH) can successfully take over language function, suggesting that early in development, the two hemispheres are largely equipotential for language (e.g., Basser, 1962; Lenneberg, 1967; Brown & Jaffe, 1975; Berl et al., 2014; Asaridou et al., 2020; cf. Bradshaw & Nettleton, 1981; Rankin et al., 1981; Vargha-Khadem et al., 1985; Pena et al., 2003; see Holland et al., 2007 and Staudt, 2007 for reviews), in spite of innate hemispheric asymmetries in anatomy (e.g., Wada et al., 1975; Chi et al., 1977; Shaw et al., 2009; Dubois et al., 2009; Glasel et al., 2011; Leroy et al., 2011; Habas et al., 2012; Leroy et al., 2015). This resilience to early brain damage has been observed for diverse kinds of brain damage, including both i) organic damage, as in cases of pre-/perinatal or early childhood stroke (e.g., Booth et al., 2000; Staudt et al., 2001, 2002; Feldman et al., 2002; Liegeois et al., 2004; Jacola et al., 2006; Newport et al., 2017; Asaridou et al., 2020; cf. Beharelle et al., 2010; see Francois et al., 2021 for a review) or epilepsy (e.g., Springer et al., 1999; Adcock et al., 2003; Brazdil et al., 2003; Woermann et al., 2003; Brazdil et al., 2005; Janszky et al., 2003, 2006; Pahs et al., 2013; see Goldman & Golby, 2005 and Hamberger & Cole, 2011 for reviews), and ii) in cases of brain injury (e.g., Vicari et al., 2000) or surgical resections (e.g., Tivarus et al., 2012), including, in some cases, of the entire left hemisphere (e.g., Basser, 1962; Boatman et al., 1999).

One important question about the development of the language network concerns the emergence and maturation of the frontal areas. In the adult brain, the frontal and temporal language areas that support high-level language comprehension appear functionally similar, showing engagement in both lexico-semantic and combinatorial semantic/syntactic processing (e.g., Keller et al., 2001; Roder et al., 2002; Fedorenko et al., 2010, 2012, 2020; Bautista & Wilson, 2016; Blank et al., 2016). However, given the protracted development of the frontal cortex (e.g., Huttenlocher, 1990; Mrzljak et al., 1990; Pfefferbaum et al., 1994; Giedd et al., 1999; Matsui et al., 2016; see Fuster, 2002 for review), frontal language areas likely develop and/or reach maturity later than the temporal ones (e.g., Ahmad et al., 2003; Wood et al., 2004; cf. Kinney et al., 1988; Pujol et al., 2006; Leroy et al., 2011), and may exhibit early functional differences.

So, how do frontal language areas emerge? There are at least two possibilities. The ***first*** is that they develop independently of the temporal language areas, albeit emerging later in life, or taking longer to reach maturity. Once these areas emerge, the white-matter fronto-temporal pathways develop and strengthen, leading to a fully mature language network (**Figure 1a**). Some evidence for this hypothesis comes from studies that have reported slow maturation of at least some intra-hemispheric pathways connecting frontal and temporal language areas. For example, Perani et al. (2011) (also Su et al., 2008; Brauer et al., 2011, 2013; Tak et al., 2016; cf. Leroy et al., 2011) have argued that the dorsal pathway (the arcuate / superior longitudinal fasciculus) does not fully mature until late childhood / early adolescence (see Friederici, 2009 for a review). In line with this evidence from DTI/DWI studies, frontal and temporal language areas in children show less synchronized activity, as assessed by functional correlations, compared to adults (e.g., Friederici et al., 2011; Xiao et al., 2016; Youssofzadeh et al., 2018; but see e.g., Satterthwaite et al., 2012 for a discussion of potential motion confounds, especially important for long-range connections). Yet, frontal language areas appear to be broadly functionally similar to those of adults already by age ∼5 (e.g., Friederici et al., 2011; Olulade 2020), arguing against the critical role of at least the dorsal pathway in their development. The ***second*** possibility is that the frontal areas emerge through the intra-hemispheric fronto-temporal pathways – perhaps through the allegedly earlier-developing ventral pathway (e.g., Brauer et al., 2013) – so the temporal language areas are critically needed to “set up” the frontal language areas (**Figure 1a**).

**Figure 1.**
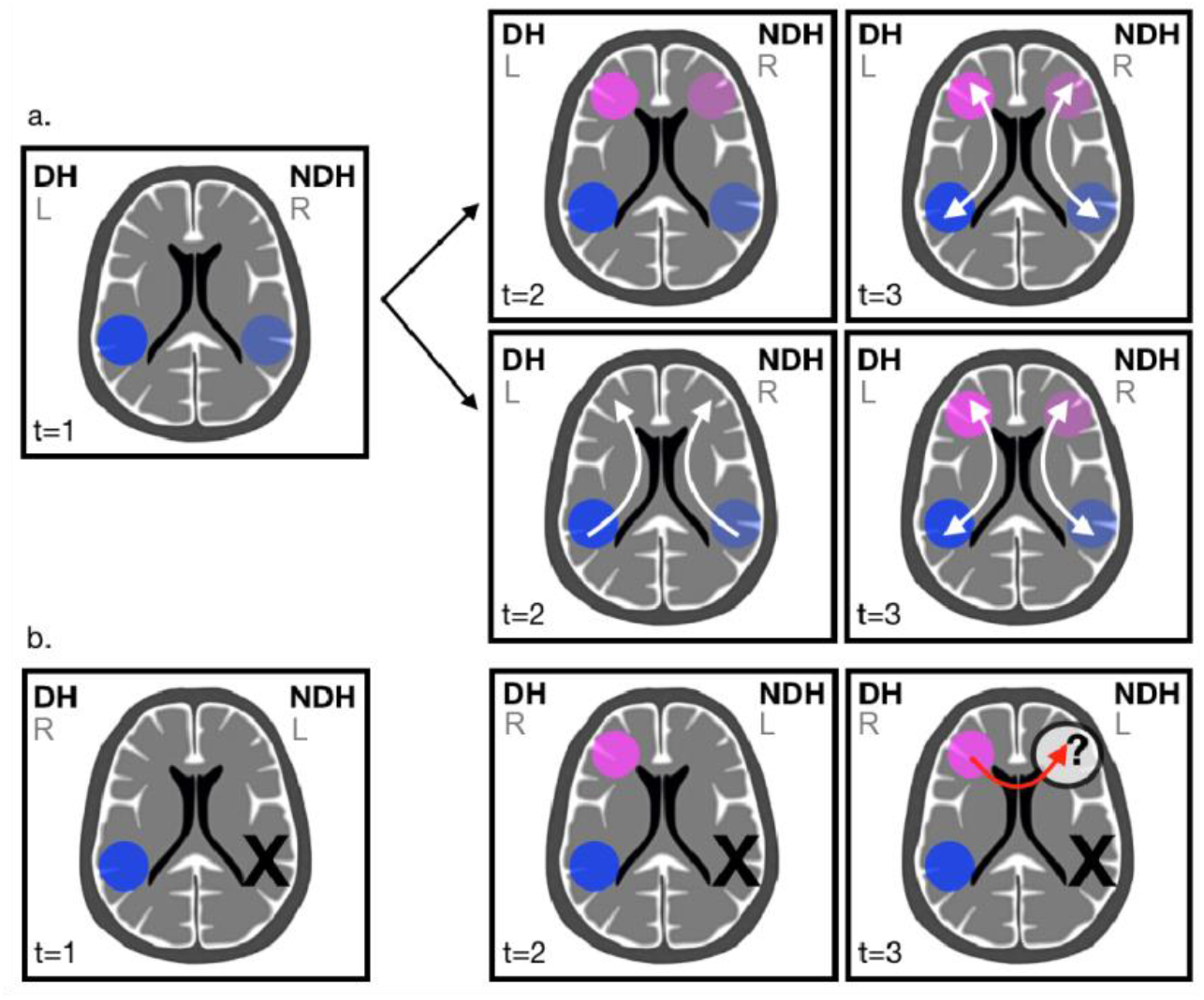
**a.** A schematic illustration of the two possibilities for how frontal language areas may develop: top – emergence *independently* of the temporal language areas, followed by the strengthening of intra-hemispheric pathways; bottom – emergence from the temporal language areas *through the intra-hemispheric connections*. In both cases, we assume that frontal language areas emerge and/or reach maturity later than temporal ones. **b.** A schematic illustration of EG’s brain with the missing left temporal lobe (marked by an X, note that the language-dominant hemisphere is now the right one, R) and the critical research question asked in the current study: namely, whether a frontal language area would develop absent the temporal lobe. Across a and b: t=1 through t=3 indicate different time points in the developmental trajectory; DH=language-dominant hemisphere (i.e., the hemisphere that eventually becomes dominant for language processing), NDH=non language-dominant hemisphere, L=left, R=right.

In either case, language areas appear to develop in both the (eventually) language-dominant hemisphere, but also in the non-dominant one, given the largely bilateral responses to language observed in childhood (e.g., Holland et al., 2001; Chou et al., 2006; Szaflarski et al., 2006; Brauer & Friederici, 2007; McNealy et al., 2011; May et al., 2011; Bonte et al., 2013; Berl et al., 2014; Olulade et al., 2020; see Holland et al., 2007 for a review). Furthermore, the language areas in the non-dominant hemisphere plausibly develop independently of those in the dominant hemisphere because bilateral language responses have been reported in individuals with agenesis of the corpus callosum (e.g., Tyszka et al., 2012; Hinkley et al., 2016), suggesting that inter-hemispheric connections, although prominent in childhood (e.g., Friederici et al., 2011; Perani et al., 2011; Naoi et al., 2013; Xiao et al., 2016; Youssofzadeh et al., 2018), are not necessary for the emergence of these areas. Instead, these inter-hemispheric connections may be used to gradually inhibit the language areas in the non-dominant hemisphere, leading to lateralized language function in adulthood (e.g., Moscovitch et al., 1976; Chiarello, 1980; Dennis, 1980; Karbe et al., 1998; Gazzaniga, 2000; Selnes, 2000; Thiel et al., 2006).

To assess the importance of the temporal language areas and tracts for the emergence of the frontal language areas, we examined language processing in an individual (EG) who was born without her left temporal lobe, likely as a result of pre-/perinatal stroke. Given early LH damage, we expected to observe a functional language network in EG’s right hemisphere. The critical question was *whether EG’s intact left lateral frontal lobe would contain language-responsive areas*. If so, that would suggest that frontal language areas can emerge without input from the ipsilateral temporal language areas. If not, that would suggest that temporal language areas are critical for the emergence of the frontal ones, and that frontal inter-hemispheric connections are not sufficient (**Figure 1b**).

## Materials and Methods

### Participants

#### Participant of interest

The participant of interest (referred throughout the manuscript as “EG”, fake initials) contacted professors in BCS, MIT in February 2016 volunteering to participate in brain research studies. EG is a highly educated right-handed female, with an advanced professional degree (four years of college-level education and three years of graduate-level education), who was aged 54 and 57 at the times of testing. EG’s parents and grandparents were stated as being right-handed, and one of her siblings as left-handed. By her own report, EG had no left temporal lobe (**Figure 2a**; see **Supplement I** for additional anatomical images). As far as she knew, this was a congenital condition; she did not suffer any head traumas or injuries as a child or adult. The lack of the left temporal lobe was discovered when an MRI scan was performed in 1987 (when EG was 25 years old and was being treated for depression). The scan was repeated in 1988, and then again in 1998, with no changes noted. Finally, the most recent clinical MRI scan (prior to testing at MIT) was conducted in 2013 when EG was suffering from headaches; no changes were observed relative the earlier scans. EG reported no problems with vision, aside from nearsightedness (corrected with glasses). With respect to speech and language, she reported no problems except for being a “terrible speller”. She had studied a foreign language (Russian) in adulthood and achieved high proficiency. We invited EG to participate in a behavioral and fMRI study at MIT. The testing took place in October 2016 (session 1) and September 2019 (session 2). The second session was conducted to ensure the robustness of the results, in line with increasing emphasis in the field on replicability (e.g., Bishop, 2019; Poldrack et al., 2017; Siegelman et al., 2019).

**Figure 2.**
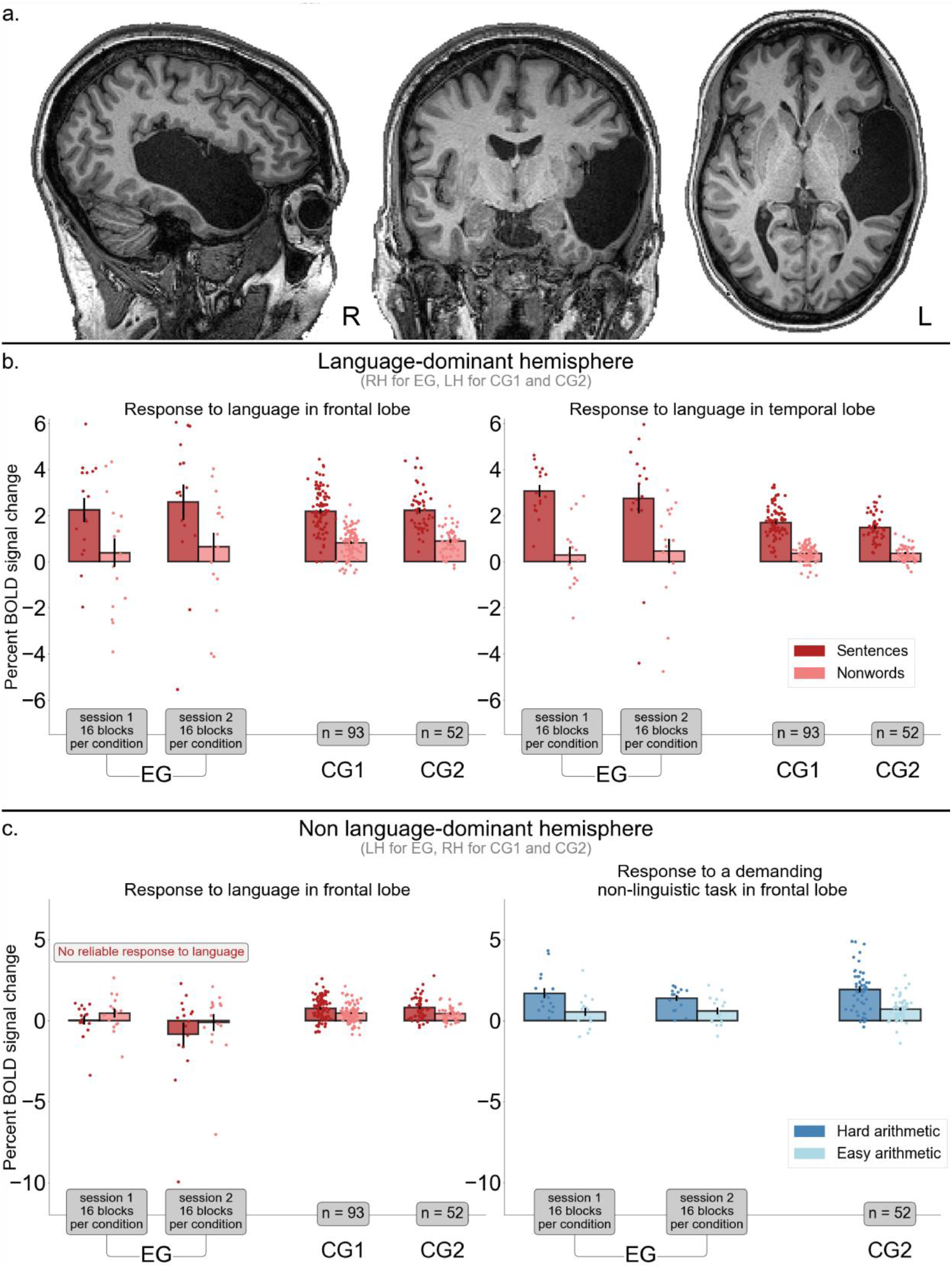
**a.** Sagittal, coronal and axial and T1-weighted images for EG (from session 1; no changes were detected in session 2). Additional lesion images are shown in **Supplement I. b.** Responses to the language localizer task in the language-dominant hemisphere (RH for EG, LH for controls): BOLD response magnitudes to sentences and nonwords. The left panel shows the responses in the frontal lobe consisting of the IFG, IFGorb, and MFG language fROIs, and the right panel shows the responses in the temporal lobe consisting of the AntTemp and PostTemp language fROIs. **c.** Responses to the language localizer task in the non language-dominant hemisphere (LH for EG, RH for controls): BOLD response magnitudes to sentences and nonwords (left panel) and the hard and easy condition of a non-linguistic arithmetic task (right panel). As in a., the frontal lobe consisted of the language fROIs in the IFG, IFGorb, and MFG (left panel), while the MD system frontal fROIs were in the PrecG, IFGop, MFG, and MFGorb (right panel). For the control groups, the error bars indicate the standard error of the mean by participants. For EG, the error bars indicate the standard error of the mean by experimental blocks.

#### Control participants

EG’s functional brain responses were evaluated relative to two controls groups. Participants for both groups were recruited from MIT and the surrounding Cambridge/ Boston, MA community and were paid for their participation. <u>Control group 1</u> (CG1) consisted of 94 native English speakers (aged 18-63 at the time of scan, mean age 25, SD 8.4; 52 females). 75 participants were right-handed (as determined by the Edinburgh handedness inventory, Oldfield 1971, or self report), 8 were left-handed/ambidextrous, and for 11, handedness information was missing. <u>Control group 2</u> (CG2) consisted of 57 native speakers of diverse languages, all proficient in English (aged 19-45 at the time of scan, mean age 28, SD 5.6; 29 females). 53 participants were right-handed, 2 were left-handed/ambidextrous, and for 2, handedness information was missing. No participants were excluded based on handedness from either group (see Willems et al. 2014, for discussion); however, we ensured that all left-handed/ambidextrous/no-handedness-information participants had a left-lateralized language network, based on the language localizer task described below. All participants in CG1 and CG2 had normal hearing and vision, and no history of language impairment. (Note that although the participants in the two control groups are, on average, younger than EG (due to availability of fMRI data for the relevant paradigms), this difference does not affect interpretation of any of the results. In fact, the age difference works against us for the main critical result, as elaborated below.)

The protocol for these studies was approved by MIT’s Committee on the Use of Humans as Experimental Subjects (COUHES). All participants gave written informed consent in accordance with the requirements of this protocol.

### Design, stimuli, and procedure

Every participant completed a language localizer task (Fedorenko et al., 2010). Because participants in CG2 were not native speakers of English, we used CG1 for the primary language comparisons. Participants in CG2 completed an arithmetic addition task that was used to localize the domain-general Multiple Demand (MD) network (e.g., Duncan, 2010, 2013; Fedorenko et al., 2013; Assem et al., 2020a). This bilateral network supports executive functions and is used here as a control, as elaborated below. EG completed a language localizer and an arithmetic addition task during each of the two visits (∼3 years apart), so in the analyses, we report the results for each of the two sessions, to establish their robustness and replicability. Each control participant completed one scanning session, and EG completed two scanning sessions during each visit. Each scanning session (for both CG participants and EG) included several additional tasks for unrelated studies and lasted approximately two hours. EG further completed a series of questionnaires and behavioral tasks, including standardized language assessments (performed during the first visit) (as detailed below).

### fMRI tasks

#### Language localizer task

The task used to localize the language network is described in detail in Fedorenko et al. (2010); the materials and scripts are available from the Fedorenko Lab website (https://evlab.mit.edu/funcloc). Briefly, we used a reading task contrasting sentences (e.g., THE SPEECH THAT THE POLITICIAN PREPARED WAS TOO LONG FOR THE MEETING) and lists of unconnected, pronounceable nonwords (e.g., LAS TUPING CUSARISTS FICK PRELL PRONT CRE POME VILLPA OLP WORNETIST CHO) in a standard blocked design with a counterbalanced condition order across runs. Each stimulus consisted of 12 words/nonwords. Stimuli were presented in the center of the screen, one word/nonword at a time, at the rate of 450ms per word/nonword. Each stimulus was preceded by a 100ms blank screen and followed by a 400ms screen showing a picture of a finger pressing a button, and a blank screen for another 100ms, for a total trial duration of 6s. Experimental blocks lasted 18s (with 3 trials per block), and fixation blocks lasted 14s. Each run (consisting of 5 fixation blocks and 16 experimental blocks) lasted 358s. Participants completed 2 runs. Participants were instructed to read attentively (silently) and press a button on the button box whenever they saw the picture of a finger pressing a button on the screen. The button-pressing task was included to help participants remain alert. The sentences > nonwords contrast targets brain regions that support lexico-semantic and combinatorial (semantic and syntactic) processing. Importantly, this localizer has been shown to robustly activate the fronto-temporal language network regardless of the specific task, materials, and modality of presentation (e.g., Fedorenko et al. 2010; Fedorenko 2014; Scott et al. 2017; Diachek, Blank, Siegelman et al., 2020; Ivanova et al., 2020). Further, the same network supports both comprehension and production (e.g., Menenti et al., 2011; Silbert et al., 2014; Hu, Small et al., in prep.).

#### Multiple Demand (MD) localizer task

The task used to localize the domain-general Multiple Demand (MD) network was an arithmetic addition task contrasting harder and easier problems in a standard blocked design with a counterbalanced condition order across runs. The easy condition involved summing two single-digit numbers whose sum could be any integer in the range of 2 to 9. The hard condition involved summing an integer in the range of 12 to 19 and an integer in the range of 2 to 9 whose sum could be any integer in the range of 21 to 28. At the end of each trial, participants were shown two numbers and performed a two-alternative forced-choice task to indicate the correct sum. The two response choices presented always differed by 2 (to prevent the participants from being able to select the correct answer using parity alone). The numbers were presented in the center of the screen for 1,450ms, followed by the response choices presented for 1,450ms and an inter-stimulus interval of 100ms. Experimental blocks lasted 15 s (with 5 trials per block), and fixation blocks lasted 15s. Each run consisted of 16 experimental blocks—8 blocks per condition—and 5 fixation blocks; a fixation block appeared at the beginning of the run and after each set of four experimental blocks, and lasted 315s. Participants completed 2 runs. The hard > easy contrast targets brain regions that support demanding cognitive tasks. This and similar contrasts between harder and easier conditions of demanding tasks have been shown to robustly activate the fronto-parietal Multiple Demand (MD) network (e.g., Fedorenko et al. 2013; Blank et al., 2014; Assem et al., 2020b; Shashidara et al., 2020).

### Behavioral tasks (EG only)

#### Language Assessment

During her first visit (in 2016), EG completed four standardized language assessment tasks: i) an electronic version of the Peabody Picture Vocabulary Test (PPVT-IV) (Dunn & Dunn, 2007); ii) an electronic version of the Test for Reception of Grammar (TROG-2) (Bishop, 2003); and iii) the Western Aphasia Battery-Revised (WAB-R) (Kertesz, 2006). PPVT-IV and TROG-2 target receptive vocabulary and grammar, respectively. In these tasks, the participant is shown sets of four pictures accompanied by a word (PPVT-IV, 72 trials) or sentence (TROG-2, 80 trials) and has to choose the picture that corresponds to the word/sentence by clicking on it. WAB-R (Kertesz, 2006) is a more general language assessment for persons with aphasia. It consists of 9 subscales, assessing 1) spontaneous speech, 2) auditory verbal comprehension, 3) repetition, 4) naming and word finding, 5) reading, 6) writing, 7) apraxia, 8) construction, visuospatial, and calculation tasks, and 9) writing and reading tasks.

#### General Cognitive Assessment

In addition to the language tasks, EG completed (also during the 2016 visit) i) an electronic version of the Kaufman Brief Intelligence Test (KBIT-2) (Kaufman & Kaufman, 2004), and ii) the 3-pictures version of the Pyramids and Palm Trees Test (Howard & Patterson, 1992). The former consists of three subtests – two verbal (Verbal Knowledge and Riddles) and one non-verbal (Matrices) – and is used to assess general fluid intelligence. The Verbal Knowledge subset consists of 60 items measuring receptive vocabulary and general information about the world; the Riddles subtest consists of 48 items measuring verbal comprehension, reasoning, and vocabulary knowledge; and the Matrices subtest consists of 46 items that involve both meaningful (people and objects) and abstract (designs and symbols) visual stimuli that require understanding of relationships among the stimuli. The Pyramids and Palm Trees test assesses non-verbal semantic cognition. The task consists of 52 trials. On each trial the participant is shown a test picture (e.g., an Egyptian pyramid) and two other pictures (e.g., a palm tree and a fur tree) and asked to choose the picture that is semantically related to the test picture (in this case, a palm tree is the correct answer). For both tests, EG’s performance was evaluated against existing norms.

### fMRI data acquisition, preprocessing, and first-level modeling

#### Data acquisition

Structural and functional data were collected on the whole-body, 3 Tesla, Siemens Trio scanner with a 32-channel head coil, at the Athinoula A. Martinos Imaging Center at the McGovern Institute for Brain Research at MIT. T1-weighted structural images were collected in 176 sagittal slices with 1mm isotropic voxels (TR=2530ms, TE=3.48ms). Functional, blood oxygenation level dependent (BOLD), data were acquired using an EPI sequence (with a 90 degree flip angle and using GRAPPA with an acceleration factor of 2), with the following acquisition parameters: thirty-one 4mm thick near-axial slices acquired in the interleaved order (with 10% distance factor), 2.1mm×2.1mm in-plane resolution, field of view of in the phase encoding (A>P) direction 200mm and matrix size 96mm×96mm, TR=2000ms and TE=30ms. Prospective acquisition correction (Thesen et al., 2000) was used to adjust the positions of the gradients based on the participant’s motion from the previous TR. The first 10s of each run were excluded to allow for steady state magnetization.

#### Preprocessing and modelling

Functional data were preprocessed and analyzed using SPM12 (using default parameters, unless specified otherwise) and supporting, custom MATLAB scripts. Preprocessing of functional data included motion correction (realignment to the mean image of the first functional run using 2^nd^-degree b-spline interpolation), direct functional normalization into a common space (Montreal Neurological Institute (MNI) template) (estimated for the mean image using trilinear interpolation), resampling into 2mm isotropic voxels, smoothing with a 4mm FWHM Gaussian filter, and high-pass filtering at 128s. For EG, SPM’s tissue probability map (TPM), standardly used in normalization was modified to include an extra layer corresponding to the lesion. Specifically, the probability for all non-lesioned tissues was set to zero, and that for the lesion volume was set to 1. A lesion mask was created in native space using automatic image segmentation, from the high-resolution structural scan and then converted into MNI space with standard normalization of the structural scan and added to the TPM.

Effects in each voxel (except those that belonged to the lesion mask) were estimated using a General Linear Model (GLM) in which each block of the experimental conditions was modeled with a boxcar function (modeling an entire block) convolved with the canonical hemodynamic response function (HRF), with nuisance regressors for linear drift removal, offline-estimated motion parameters, and outlier time points (where the scan-to-scan differences in global BOLD signal were above 5 standard deviations, or where the scan-to-scan motion was above 0.9mm). We modeled the experimental blocks separately in order to be able to estimate variance for each condition for EG.

### Definition of language and MD functional regions of interest (fROIs)

Responses to each block of each condition were extracted from regions of interest that were defined functionally in each individual participant (e.g., Saxe, et al., 2006; Fedorenko, et al., 2010; Nieto-Castañón and Fedorenko, 2012; Fedorenko, 2021). Two sets of functional regions of interest (fROIs) were defined—one for the language network and one for the MD network. In particular, fROIs were constrained to fall within a set of ‘masks’ which delineated the expected gross locations of activations for the relevant contrast and were sufficiently large to encompass the extent of variability in the locations of individual activations. (This inter-individual topographic variability, well-documented in prior work (e.g., Fedorenko et al., 2010, Mahowald & Fedorenko, 2016, Braga et al., 2020, and Affourtit et al., in prep.; see Fedorenko & Kanwisher, 2009 and Fedorenko & Blank, 2020 for reviews) is the key motivation for the development of paradigms that robustly identify the areas and networks of interest at the individual-subject level (Kanwisher et al., 1997; Saxe et al., 2006; Fedorenko et al., 2010; Fedorenko, 2021).) These masks were derived from probabilistic activation overlap maps in independent sets of participants, as described in Fedorenko et al. (2010), and elaborated below.

#### Language fROIs

To define the language fROIs, each individual map for the *sentences > nonwords* contrast from the language localizer was intersected with a set of five binary masks. These masks were derived from a probabilistic activation overlap map for the language localizer contrast in 220 participants. Five language fROIs were defined in the dominant hemisphere (RH for EG, and LH for the controls): three on the lateral surface of the frontal cortex (in the inferior frontal gyrus, IFG, and its orbital part, IFGorb, as well as in the middle frontal gyrus, MFG), and two on the lateral surface of the temporal and parietal cortex (in the anterior temporal cortex, AntTemp, and posterior temporal cortex, PostTemp). Further, in the controls, five homotopic fROIs were defined in the non language-dominant (right) hemisphere; and in EG, three homotopic fROIs were defined in the frontal lobe of the non language-dominant (left) hemisphere. Following prior work (e.g., Blank et al., 2014), to define the RH fROIs, the LH language masks were transposed onto the RH, allowing the LH and RH fROIs to differ in their precise locations within the masks. All masks are available for download from https://evlab.mit.edu/funcloc/.

#### MD fROIs

To define the MD fROIs, each individual map for the *hard > easy arithmetic* contrast was intersected with a set of eight binary masks in the frontal cortex (we do not here examine the parietal MD fROIs because our focus is on the frontal lobe). These masks were derived from a probabilistic activation overlap map for a similar contrast (based on a working memory task; see Fedorenko et al., 2013 for evidence that these contrasts activate the same network) in 197 participants. These masks covered the frontal components of the fronto-parietal MD network and closely overlapped with a set of anatomical masks used in Fedorenko et al. (2013). Four fROIs in the frontal cortex were defined in both EG and the controls (in the precentral gyrus, LH/RH PrecG, the opercular part of the inferior frontal gyrus, LH/RH IFGop, the middle frontal gyrus, LH/RH MFG, and its orbital part, LH/RH MFGorb).

Independent data were used to define the regions of interest and extract blockwise responses. In particular, condition-level contrasts (averaging across blocks) were defined with one run and responses for each block were extracted from the other run. For example, the contrast *sentences blocks in run 1 > nonwords blocks in run 1* was used to define the fROIs whose responses to the blocks in run 2 were estimated. This procedure was then repeated for the other run, defining fROIs with run 2 to estimate block-wise responses in run 1. A given fROI was defined as the top 10% of most localizer-responsive voxels within each mask.

### Critical fMRI analyses

A series of analyses were performed to address three key research questions. First, we asked whether EG’s language network in the right hemisphere (her language-dominant hemisphere) is comparable to the language network in the left hemisphere of the control participants. Next and critically, we asked whether EG’s left frontal lobe contains language-responsive areas. And finally, we assessed the general functionality of EG’s left frontal lobe by probing its responses to the arithmetic addition task, which has been previously shown to robustly activate the bilateral MD network. We describe the specific analyses, organized by these questions, below. To test for statistical significance in all analyses, we exploit two statistical methods: i) A Bayesian assessment of the atypicality of a single-case (EG) score against a set of control scores (Crawford & Garthwaite, 2007), and ii) Linear mixed-effects models (Barr et al., 2013). Analyses of the language network used CG1 as the primary control group (for completeness, we show that the results generalize to CG2), whereas analyses of the MD network only used CG2 (because the arithmetic task was not included in CG1 participants). Each statistics test was performed on the data from each of the two sessions, to ensure robustness and replicability.

*1. Is EG’s RH language network similar to the LH language network in control participants?*

To compare EG’s language-dominant hemisphere language network to that of controls, we compared the responses to the language task in EG’s RH frontal and temporal language fROIs to those in the controls’ LH frontal and temporal fROIs (**Figure 2b**).

*2. Does EG’s LH frontal lobe support language processing?*

To test for the presence of language responses in EG’s LH frontal lobe, we compared the responses to the language task in EG’s LH frontal language fROIs to those in the controls’ RH frontal fROIs (**Figure 2c, left panel**).

*3. Does EG’s LH frontal lobe support non-linguistic processing?*

Finally, we performed an analysis to ensure that EG’s LH frontal lobe is functional even if it does not support language processing. To do so, we examined her LH frontal responses to a non-linguistic task—arithmetic processing. We compared the strength of activation between EG’s non language-dominant frontal lobe, and the non language-dominant frontal lobe of CG2 participants (**Figure 2c, right panel**).

#### Statistical tests

For each of these analyses, we compared EG’s effect size value to the distribution of the corresponding effect size values in the control participants using two statistical approaches: 1) A Bayesian test for single-case assessment (Crawford & Garthwaite, 2007) using the *psycho* (Makowski, 2018) package in R (10,000 iterations), and 2) Linear mixed effects (LME) regression models (Barr et al., 2013) implemented in R. With respect to the first approach, it has been suggested that exploiting case-control comparisons using z-score-based methods is appropriate given large control samples because the sampling distribution of the statistic of interest will be approximately normal (McIntosh and Rittmo, 2021). The minimum sample size is often context-dependent, but has been suggested to be unproblematic for sample sizes of n >= 50 (Crawford and Howell, 1998). The Crawford *p*-values reported are two-tailed: they provide estimates of the probability that a member of the control population would exhibit a larger difference in either direction (Crawford and Garthwaite, 2007).

With respect to the second approach, the LME models were implemented using the *lmer* function from the *lme4* R package (Bates et al., 2015) and statistical significance of the model effects was evaluated using the *lmerTest* package (Kuznetsova et al., 2017) with Satterthwaite’s method for approximating degrees of freedom. Effect sizes obtained for each experimental block were modelled with a fixed effect for group (EG vs. controls), condition (language: *sentences* vs. *nonwords* condition; MD: *hard* vs. *easy arithmetic* condition), and an interaction between group and condition. To account for unexplained differences between participants, fROIs, and experimental blocks, the model additionally included random intercepts by participant, fROI, and experimental block (the fROIs were further grouped by hemisphere and lobe; for the language network analyses, we examined language-dominant hemisphere frontal fROIs, language-dominant hemisphere temporal fROIs, and non language-dominant hemisphere frontal fROIs, and for the MD network, we examined LH frontal fROIs). The models were coded with the control group in the intercept. For the language LME models, the *nonwords* condition was modelled in the intercept. For the MD LME models, the *easy arithmetic* condition was modelled in the intercept. The models were fitted using maximum likelihood estimation. The R^2^ of the models was determined using GLMM in R (Nagakawa et al., 2013; Johnson, 2014; Nagakawa et al., 2017).

To test our critical question of whether EG significantly differs from the control population, we performed additional hypothesis testing using likelihood ratio tests (LRT) by comparing the full LME model (see **Supplement II**) to an ablated LME model (see **Supplement II**) without the critical interaction term. The interaction term consisted of an interaction between EG and the controls. The null hypotheses *H*_0_ is that the likelihoods of the two models are equivalent. Thus, if *H*_0_ is rejected, the observed response cannot be explained by the ablated LME model without the interaction term, and thus EG differs from the control distribution. The Chi Square value, X^2^, was used as the test statistic and was implemented using the *ANOVA* function from the *lme4* package.

For each LME model reported, we provide (in **Supplement II**) a table with model formulae, effect size estimates, standard error estimates, *t*-statistics, degrees of freedom, *p*-values, and R^2^ values (the conditional R^2^ is reported, which accounts for the variance explained by the entire model, including both fixed and random effects). For each Crawford test reported, we provide (in **Supplement III**) a table with effect sizes, percentiles, credible intervals and control group mean/standard deviation.

For the Crawford analyses, a test was performed for each fROI group by averaging across the block-wise fROI effect sizes (to minimize multiple comparisons given that the Crawford test is a Bayesian alternative to a *t*-test). For the LME analyses, an LME model was fitted for each fROI group while still modeling individual fROIs as random effects as noted above (see **Supplement II** for exact model formulae).

Prior to statistical modeling, participants with a mean response across conditions and blocks in the 0.1th and 99.9th percentile of the data within each fROI group were excluded from all analyses. This resulted in the exclusion of one participant from CG1, leaving us with n=93 participants, and five participants from CG2, leaving us with n=52 participants.

## Results

### Behavioral results (EG only)

#### Language Assessment

In line with EG’s self-report, she performed within normal range on all language assessment tasks. She got 90% correct on PPVT, 99% correct on TROG, and 97.6, 98.6, and 98.4 on the aphasia, language, and cortical quotients of the WAR-B (the criterion cut-off score for diagnosis of aphasia is an aphasia quotient of 93.8). EG’s performance was therefore not distinguishable from the performance of neurotypical controls.

#### General Cognitive Assessment

EG performed within normal range on both general cognitive assessments. Her KBIT scores were 130 (98^th^ percentile) on the verbal composite assessment (across the two subtasks; see *Methods*), 54 (79^th^ percentile) on the non-verbal assessment, and 122 (93^rd^ percentile) overall composite assessment. She answered 51 of the 52 questions correct on the Pyramids and Palm Trees task.

### fMRI results

This section is organized in terms of the three questions laid out in *Methods/Critical fMRI analyses*. As noted above, in line with increasing emphasis on robustness and replicability (e.g., Bishop, 2019; Poldrack et al., 2017; Siegelman et al., 2019), we tested EG twice (3 years apart). The results are reported as session 1 and session 2 in the sections below.

#### 1. Is EG’s RH language network similar to the LH language network in control participants?

The LME model showed a significant effect of condition for the language-dominant hemisphere frontal language fROIs (sentences > nonwords, session 1: β=1.362, p<.001; session 2, β=1.362, p<.001) and for the language-dominant hemisphere temporal language fROIs (sentences > nonwords, session 1: β=1.261, p<.001; session 2, β=1.261, p<.001). No significant effect of group was observed either for the frontal fROIs (EG > controls, session 1: β=-0.454, p=0.527; session 2, β=-0.2, p=0.781) or for the temporal language fROIs (EG > controls, session 1: β=- 0.071, p=0.876; session 2, β=0.107, p=0.814). No significant interaction between condition and group was observed for the frontal language fROIs (session 1: X^2^(1)=1.392, p=0.238; session 2: X^2^(1)= 1.906, p=0.167), but EG showed significantly stronger responses to sentences compared to the control participants in the temporal language fROIs, as evidenced by a significant condition by group interaction (session 1: X^2^(1)=27, p<.001; session 2: X^2^(1)=11.747, p<.001). The Crawford test yielded similar results: EG did not differ significantly from the control participants in the frontal language fROIs (session 1: p=0.237; session 2: p=0.201), but showed significantly stronger responses in the temporal language fROIs (session 1: p=0.003; session 2: p=0.031).

#### 2. Does EG’s LH frontal lobe support language processing?

The LME model showed a significant effect of condition for the non language-dominant hemisphere frontal language fROIs (sentences > nonwords, session 1: β=0.33, p=0.008; session 2: β=0.33, p=0.009). No significant effect of group was observed (EG > controls, session 1: β=0.017, p=0.977; session 2: β=-0.555, p=0.337). Critically, however, a significant interaction between condition and group was observed (session 1: X^2^(1)=4.925, p=0.026; session 2: X^2^(1)=8.886, p=0.003). As can be clearly seen in **Figures 2b and 3a**, EG did not show a reliable sentence > nonwords in either session, even numerically (in fact, in both sessions, the nonwords condition was numerically higher than the sentence condition). The Crawford test yielded similar results: EG differed significantly from control participants in the prefrontal non language-dominant hemisphere (session 1: p=0.053; session 2: p=0.015). Thus, these results demonstrate that EG had significantly lower activation to sentences compared to control participants in the prefrontal non language-dominant hemisphere. (It is worth noting that the fact that the control participants are younger than EG does not bias the results / affect their interpretation. In general, more bilateral language responses have been reported in aging brains (e.g. Logan et al., 2002; Tyler et al., 2010; see Diaz et al., 2016 for a review); so if our control participants were older, the difference with EG (who shows no response in the non language-dominant hemisphere) would likely be even larger and more statistically pronounced.)

**Figure 3.**
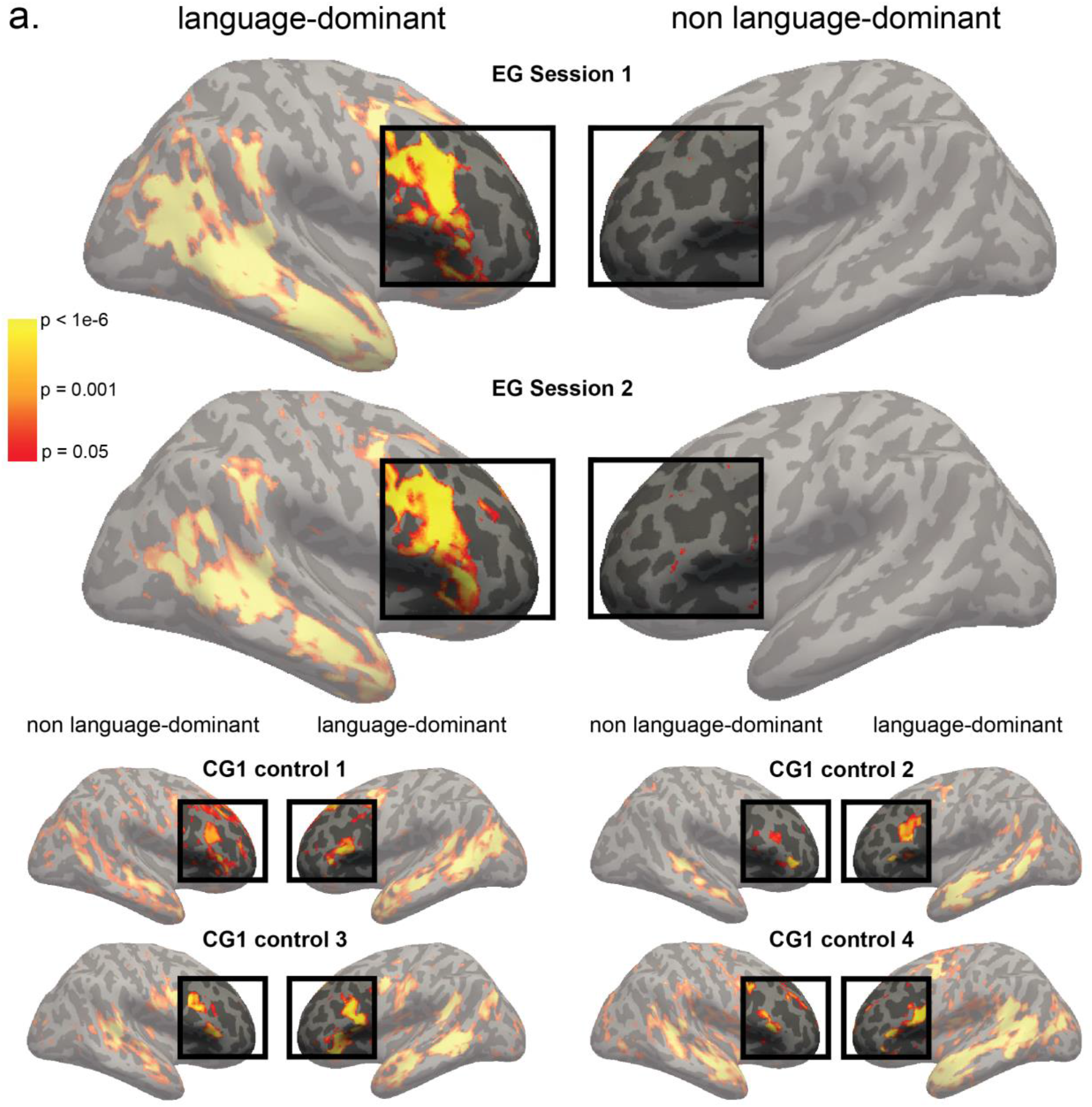
**a.** Surface projection of language activation maps (sentences > nonwords contrast) for EG and several of the control participants. The data were analyzed in the volume space and co-registered and resliced using SPM12 to Freesurfer’s standard brain CVS35 (combined volumetric and surface-based (CVS)) in the MNI152 space using 4th degree B-spline interpolation. The maps for each participant were projected onto the cortical surface using mri_vol2surf in Freesurfer v6.0.0 with a projection distance of 1.5mm. The surface projections were visualized on an inflated brain in the MNI152 space. The upper panel shows EG’s brain (language-dominant: RH, non language dominant: LH) for sessions 1 and 2 (conducted three years apart). The lower panel shows several representative participants from CG1 (language-dominant: LH, non language-dominant: RH). The figures illustrate the fact that control participants, but not EG, show frontal responses during language processing in their non language-dominant hemisphere, in line with what the statistical tests reveal.

**Figure 3.**
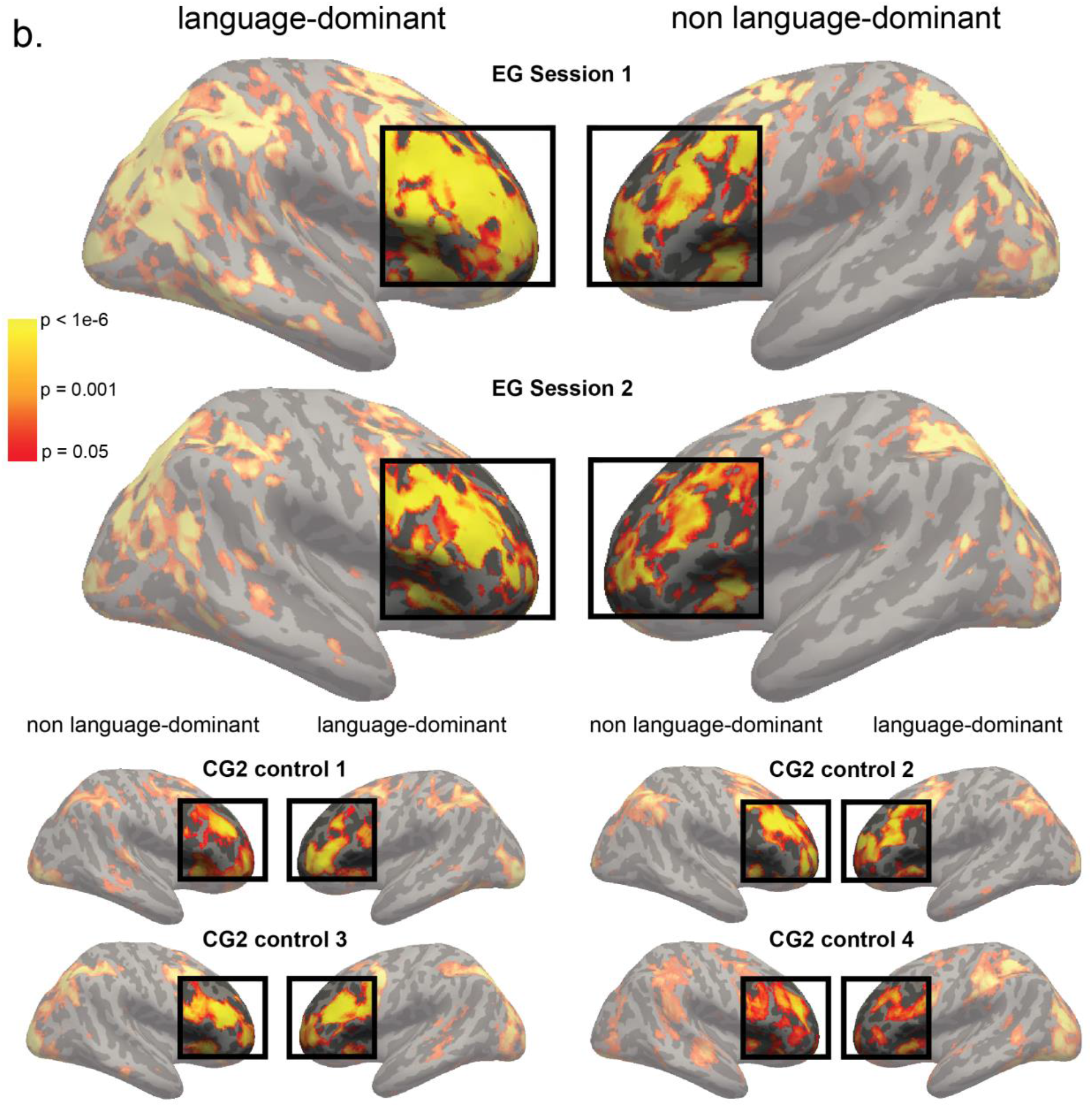
**b.** Surface projection of MD activation maps (hard > easy arithmetic contrast) for EG and several of the control participants. Same procedure as in Figure 3.a. The figures illustrate the fact that, like the control participants, EG show robust frontal responses during a demanding cognitive task in her left frontal cortex, which suggests that this part of the cortex is perfectly functional and capable of supporting some high-level cognitive functions.

#### 3. Does EG’s LH frontal lobe support non-linguistic processing in EG?

To rule out the possibility that EG’s LH frontal lobe does not respond to any high-level cognitive tasks, we examined responses to an arithmetic addition task, which robustly activates the domain-general Multiple Demand (MD) network (e.g., Fedorenko et al., 2013; Amalric & Dehaene, 2019). The MD network consists of a network of bilateral frontal and parietal brain areas (e.g., Duncan, 2010, 2013; Fedorenko et al., 2013; Assem et al., 2020a), and so has a strong presence in the LH frontal lobe. The LME model showed a significant effect of condition for LH frontal MD fROIs (hard > easy, session 1: β=1.194, p<.001; session 2: β=1.194, p<.001). No significant effect of group was observed (EG > controls, session 1: β=-0.167, p=0.863; session 2: β=-0.113, p=0.907), and no significant interaction effect between condition and group (session 1: X^2^(1)=0.02, p=0.888; session 2: X^2^(1)=1.095, p=0.295) (see **Figure 3b**). The Crawford test yielded similar results: EG did not differ significantly from the control participants in the left frontal MD fROIs (session 1; p=0.453, session 2; p=0.335).

## Discussion

In the current study, we investigated whether frontal language areas emerge absent the ipsilateral temporal language areas. We examined language processing in the brain of an individual (EG) lacking her LH temporal lobe (likely due to a pre-/perinatal stroke). In line with past work on individuals with early left-hemisphere damage / removal (e.g., Booth et al., 2000; Staudt et al., 2001; Jacola et al., 2006; Newport et al., 2017; Asaridou et al., 2020; Vicari et al., 2000; Basser, 1962; Boatman et al., 1999), EG exhibited a functional language network in her RH, and her linguistic abilities were within normal range. In fact, her verbal IQ was in the 98th percentile. However, we found no evidence of language-responsive areas in EG’s LH frontal lobe, in contrast to a large control group, who show robust frontal responses to language in the non language-dominant hemisphere. Another network supporting high-level cognitive functions—the Multiple Demand (MD) network—was robustly present in EG’s left frontal lobe (similar to controls), suggesting that the cortex in this part of the brain is perfectly functional in general and capable of supporting high-level cognition. We take these results to suggest that frontal language areas do not emerge without the ipsilateral temporal language areas. In the remainder of the Discussion, we discuss a few issues that this study informs or raises.

### Frontal language areas do not emerge in the absence of temporal language areas

The critical question we asked is whether frontal language areas would emerge in a brain that is lacking the ipsilateral temporal lobe. We laid out two possibilities for how frontal language areas may emerge. The *first* is that they develop independently of the temporal language areas, in which case the absence of the temporal lobe should not matter. And the *second* is that the frontal areas emerge through the intra-hemispheric fronto-temporal pathways from the temporal language areas, which likely emerge earlier because of their proximity to the speech-responsive auditory cortex (Norman-Haignere et al., 2015; Overath et al., 2015). Based on past work, it was already known that in childhood, language areas appear to develop bilaterally (e.g., Holland et al., 2001; Chou et al., 2006; Szaflarski et al., 2006; Brauer & Friederici, 2007; McNealy et al., 2011; May et al., 2011; Bonte et al., 2013; Berl et al., 2014; Olulade et al., 2020; see Holland et al., 2007 for a review) and independently in each hemisphere, as evidenced by bilateral language responses in individuals with agenesis of the corpus callosum (e.g., Tyszka et al., 2012; Hinkley et al., 2016). EG’s data further inform the development of the language system by showing that the temporal language areas and the intra-hemispheric fronto-temporal pathways appear to be critically needed to “set up” the frontal language areas. Given that the ventral fronto-temporal pathway, which runs through the extreme capsule (EmC) and external capsule (EC) through the inferior fronto-occipital fasciculus (IFOF), appears to mature early, being already detectable at birth (e.g., Brauer et al., 2013), we speculate that this is the pathway that supports the development of the frontal language areas.

### One hemisphere is sufficient to implement the language system

EG adds to the body of work that has suggested that a single hemisphere is perfectly sufficient to support language comprehension and production (e.g., Basser, 1962; Lenneberg, 1967; Brown & Jaffe, 1975; Berl et al., 2014; Asaridou et al., 2020). We found robust responses to language comprehension in both RH frontal and temporal language areas. The magnitude of response in EG’s RH frontal language areas was similar to that in the control participants’ language-dominant hemisphere frontal areas. The magnitude of response in EG’s RH temporal language areas was higher compared to the controls. Whether this higher magnitude of response is compensatory is difficult to determine. The mean of EG’s temporal language areas’ response magnitude overlaps with the distribution of the controls’ magnitudes, so it clearly not impossible for neurotypical individuals to exhibit this level of response. Relatedly, Asaridou et al. (2020) recently reported a case of a 14-year old child born without the left hemisphere. For a language comprehension task, they observed activation patterns in right frontal and temporal brain regions that were similar to those in age-matched neurotypical children (although the strength of the response was not directly compared, only the general topography). However, they found that the dorsal white matter tracts (the direct and anterior segments of the arcuate fasciculus) that connect areas active during language processing were larger in their participant of interest compared to a control population. Asaridou et al. suggested that these stronger dorsal tracts may play a compensatory role by providing faster and more reliable transfer of information between frontal and temporal language areas. However, whether stronger temporal lobe responses and/or larger dorsal tracts constitute ubiquitous features of brains with early extensive LH damage remains to be discovered.

An important aspect of EG’s case, as well as the case reported in Asaridou et al. (2020) and cases of early LH hemispherectomies (e.g., Basser, 1962; Boatman et al., 1999; see Lidzba et al., 2021 for a review), is that not only a single hemisphere appears to be sufficient, but the *right* hemisphere—i.e., the non language-dominant hemisphere in most neurotypical adults—appears to be perfectly suitable for language function. Although some have argued that the LH may be especially well-suited for language at birth (e.g., Bradshaw & Nettleton, 1981; Rankin et al., 1981; Vargha-Khadem et al., 1985; Pena et al., 2003), evidence continues to accumulate for the equipotentiality of the two hemispheres for language. Olulade et al. (2020) recently argued that bilateral—and presumably redundant—representation and processing of language in early childhood make the language system robust to early damage, so that even severe damage to, or even complete removal of, one hemisphere leaves a functional language system in place. That said, many questions remain about the role of the two hemispheres in language. For example, why does language end up in the LH in most individuals (e.g., Corballis, 2009; Ocklenburg et al., 2013; Sha et al., 2021)? Why doesn’t language remain bilaterally and redundantly represented and processed throughout life, which would be hugely advantageous for protection from late brain damage (e.g., see Vallortigara & Rogers, 2005, for a general discussion of the advantages of hemispheric dominance)? And how is linguistic labor distributed between the LH and RH language networks in adults?

### The organization of EG’s LH frontal lobe

Given that EG’s LH frontal lobe appears to not contribute to language processing, what perceptual, motor, or cognitive functions do the areas that would belong to the language network in neurotypical adults support in EG? We don’t have an answer yet. What we report in the current study is that components of the domain-general MD network are robustly present in EG’s left frontal lobe and show similar magnitudes of response to those in the control participants. Similarly, as can be seen in **Figure 3b**, the general topography of activation for the MD localizer task looks similar to what has been previously reported for NT individuals (e.g., Fedorenko et al., 2013; Shashidara et al., 2019) and what is observed in the current control group, albeit perhaps more extensive. This result suggests that at least some of the organization of EG’s left frontal lobe is intact. But whether/how the would-be language areas are repurposed remains to be discovered.

One interesting point worth making is that we here used an arithmetic addition task as our MD localizer task. In NT individuals, math and language draw on distinct networks (e.g., Varley et al., 2005; Fedorenko et al., 2011; Monti et al., 2012; Amalric & Dehaene, 2019) but i) math processing is generally left-lateralized (Monti et al., 2012; Amalric & Dehaene, 2019), and ii) language and math processing tend to co-lateralize (Pinel & Dehaene, 2010). It is therefore interesting that we see robust responses to an arithmetic task in EG’s left frontal lobe in spite of the fact that no language responses are detected there.

### The general decline of single-case studies

A methodological point is also worth making. The number of published neuroscience papers on single-case studies is steadily declining (e.g., Streese & Tranel, 2021, Medina and Fischer-Baum, 2017; Fellows et al., 2005). However, from the earliest days of cognitive neuroscience (e.g., Broca, 1861), such studies have provided critical insights into the architecture of the human mind and brain (e.g., Caramazza, 1986; Caramazza & Coltheart, 2006). As Streese & Tranel (2021) suggest, combining behavioral and fMRI approaches in the study of unusual brains can be especially powerful, including informing hypotheses that cannot be tested in neurotypical individuals. For example, the question we tackled in the current paper—whether frontal language areas emerge independently of temporal areas—simply cannot be answered without turning to atypical brains, and EG’s brain had just the right properties to ask and answer it. Of course, generalizing the current findings to other cases similar to EG would certainly be useful, but ‘deep data’ style investigations on one or a few individuals, like the one carried out here, are not to be underestimated (e.g., see Gratton & Braga, 2021, along with the other articles in the special issue on deep imaging for extensive discussions of the importance of careful investigations where ample data are collected within each individual). In the current study, we a) used extensively validated paradigms that have been shown to elicit reliable responses within individuals (in hundreds of participants across dozens of past studies), b) directly replicated the critical findings across two testing sessions three years apart, and c) performed careful statistical comparisons (using several analytic approaches) of individual-level neural markers (previously established to be reliable within individuals; Mahowald & Fedorenko, 2016) between the participant of interest and large control groups. Given the rise, over the last decade, of ‘deep neuroscience’ approaches in brain imaging work on neurotypical individuals (e.g., DiNicola & Braga, 2021; Gratton & Braga, 2021; Fedorenko, 2021; Naselaris et al., 2021; Noble et al., 2021; Poldrack, 2021; Smith et al., 2021), we hope that rigorous case studies of atypical brains will also make a comeback.

### Limitations of scope

Aside from the limitations inherent in the single-case study approach, we have here focused on the high-level language network, i.e., brain regions that support the processing of word meanings and combinatorial semantic/syntactic processing (e.g., Fedorenko et al. 2012b, 2020; Bautista and Wilson 2016; Blank et al. 2016). In the future, we plan to additionally examine EG frontal lobe’s response to lower-level speech perception and speech articulation tasks. We expect that those functions would be concordant with what we found for higher-level language areas, but it remains to be established empirically. We also plan to further investigate the organization of EG’s left frontal lobe in an effort to understand what functions the areas that would typically perform language processing support in her brain. Finally, we have so far focused on the cortical language responses. A recent review on brain plasticity supporting language recovery after perinatal stroke (Francois et al., 2021) found that good language outcomes (besides the reorganization of language processing to the right hemisphere) were associated with increased activity in the left cerebellum. Future work should probe cerebellar and sub-cortical language responses to paint a more complete picture of language processing in atypical brains.

## Acknowledgements

We would like to thank our participant – EG – for her time and patience. We would also like to acknowledge the Athinoula A. Martinos Imaging Center at the McGovern Institute for Brain Research at MIT, and its support team (Steve Shannon, and Atsushi Takahashi), Saima Malik-Moraleda and Dima Ayyash for help in collecting the data for the second control group, and Anne Billot and Swathi Kiran for help in evaluating EG’s linguistic abilities. EF was supported by the NIH awards R00-HD057522, R01-DC016607, and R01-DC016950, by a grant from the Simons Foundation to the Simons Center for the Social Brain at MIT, and by research funds from the Department of Brain and Cognitive Sciences, and the McGovern Institute for Brain Research.

## Notes

### Competing Interest Statement

The authors have declared no competing interest.

