## Supplementary Information for "Frontal language areas do not emerge in the absence of temporal language areas: A case study of an individual born without a left temporal lobe"

### Supplemental Information I

**Sagittal (right to left)** **Coronal (posterior to anterior)** **Axial (inferior to superior)**

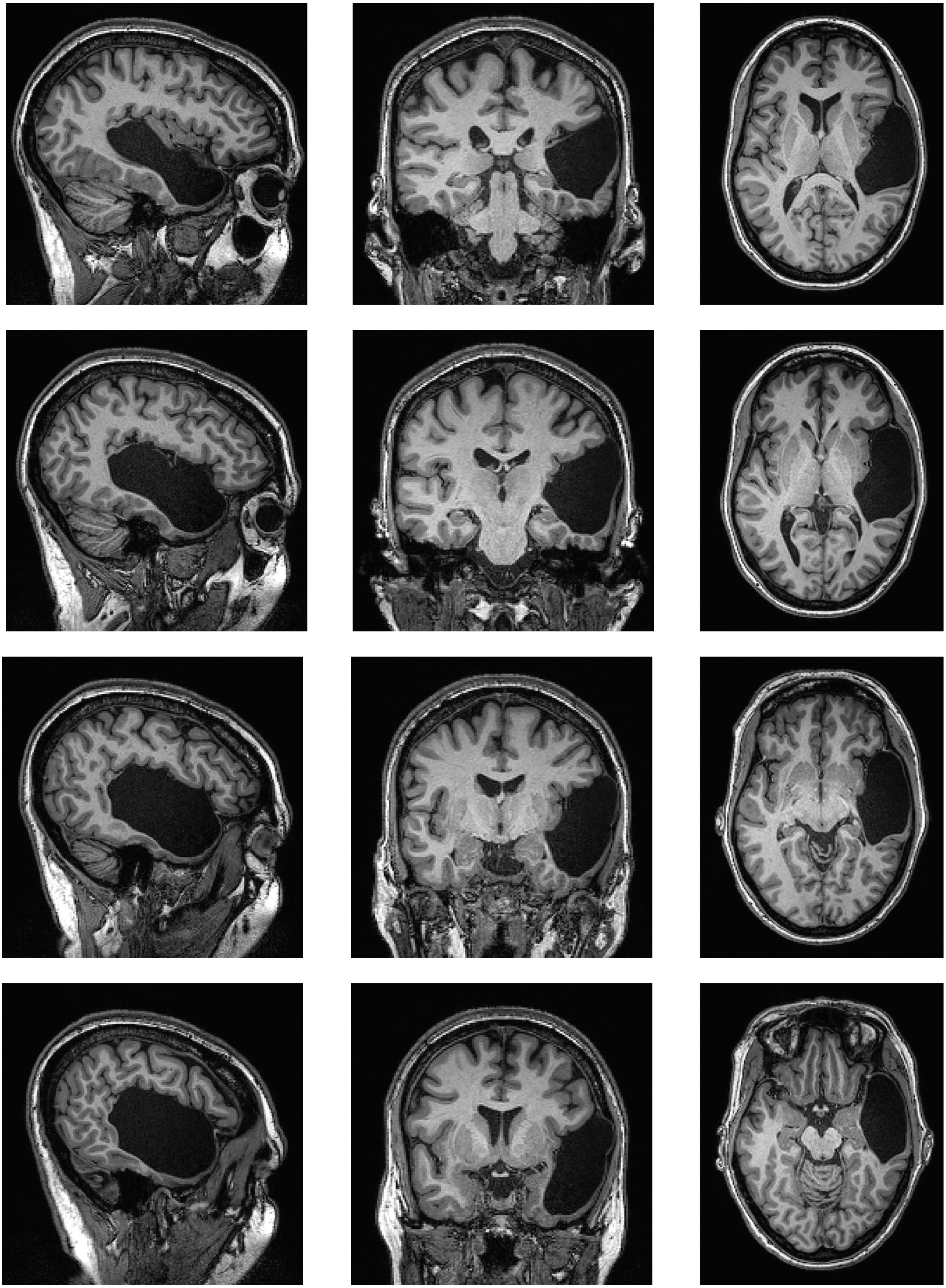

### Supplemental Information II

$$\text{Full model} = \mathit{effectsize} \sim 1 + \mathit{group} + \mathit{condition} + \mathit{group} * \mathit{condition} + (1|\mathit{participant}) + (1|froi) + (1|\mathit{block})$$

$$\text{Ablated model} = \mathit{effectsize} \sim 1 + \mathit{group} + \mathit{condition} + (1|\mathit{participant}) + (1|froi) + (1|\mathit{block})$$

Table 1: Full model, Fixed effects coefficients, language network, FDM, EG Session 1, R-Squared = 0.2428

| | Estimate | Std. Error | df | t value | Pr(>\|t\|) |
|---|---|---|---|---|---|
| (Intercept) | 0.85 | 0.30 | 4.47 | 2.79 | 0.04 |
| condS | 1.36 | 0.17 | 31.18 | 8.03 | 0.00 |
| groupEG1 | -0.45 | 0.72 | 173.24 | -0.63 | 0.53 |
| condS:groupEG1 | 0.49 | 0.42 | 13836.99 | 1.18 | 0.24 |

Table 2: Ablated model, Fixed effects coefficients, language network, FDM, EG Session 1, R-Squared = 0.2427

| | Estimate | Std. Error | df | t value | Pr(>\|t\|) |
|---|---|---|---|---|---|
| (Intercept) | 0.84 | 0.30 | 4.47 | 2.78 | 0.04 |
| condS | 1.36 | 0.17 | 31.16 | 8.05 | 0.00 |
| groupEG1 | -0.21 | 0.69 | 145.07 | -0.30 | 0.76 |

Table 3: Full model, Fixed effects coefficients, language network, FDM, EG Session 2, R-Squared = 0.2394

| | Estimate | Std. Error | df | t value | Pr(>\|t\|) |
|---|---|---|---|---|---|
| (Intercept) | 0.85 | 0.30 | 4.44 | 2.81 | 0.04 |
| condS | 1.36 | 0.17 | 31.17 | 8.20 | 0.00 |
| groupEG2 | -0.20 | 0.72 | 173.51 | -0.28 | 0.78 |
| condS:groupEG2 | 0.58 | 0.42 | 13836.99 | 1.38 | 0.17 |

Table 4: Ablated model, Fixed effects coefficients, language network, FDM, EG Session 2, R-Squared = 0.2393

| | Estimate | Std. Error | df | t value | Pr(>\|t\|) |
|---|---|---|---|---|---|
| (Intercept) | 0.84 | 0.30 | 4.44 | 2.81 | 0.04 |
| condS | 1.37 | 0.17 | 31.16 | 8.23 | 0.00 |
| groupEG2 | 0.09 | 0.69 | 145.08 | 0.13 | 0.90 |

Table 5: Full model, Fixed effects coefficients, language network, TDM, EG Session 1, R-Squared = 0.3393

| | Estimate | Std. Error | df | t value | Pr(>\|t\|) |
|---|---|---|---|---|---|
| (Intercept) | 0.36 | 0.17 | 2.97 | 2.08 | 0.13 |
| condS | 1.26 | 0.10 | 31.18 | 13.15 | 0.00 |
| groupEG1 | -0.07 | 0.45 | 180.35 | -0.16 | 0.88 |
| condS:groupEG1 | 1.51 | 0.29 | 9165.98 | 5.20 | 0.00 |

Table 6: Ablated model, Fixed effects coefficients, language network, TDM, EG Session 1, R-Squared = 0.3374

| | Estimate | Std. Error | df | t value | Pr(>\|t\|) |
|---|---|---|---|---|---|
| (Intercept) | 0.36 | 0.17 | 2.97 | 2.05 | 0.13 |
| condS | 1.27 | 0.10 | 31.15 | 13.26 | 0.00 |
| groupEG1 | 0.69 | 0.43 | 145.09 | 1.60 | 0.11 |

Table 7: Full model, Fixed effects coefficients, language network, TDM, EG Session 2, R-Squared = 0.3318

| | Estimate | Std. Error | df | t value | Pr(>\|t\|) |
|---|---|---|---|---|---|
| (Intercept) | 0.36 | 0.18 | 2.90 | 2.07 | 0.13 |
| condS | 1.26 | 0.09 | 31.17 | 13.52 | 0.00 |
| groupEG2 | 0.11 | 0.45 | 181.15 | 0.24 | 0.81 |
| condS:groupEG2 | 1.01 | 0.29 | 9165.98 | 3.43 | 0.00 |

Table 8: Ablated model, Fixed effects coefficients, language network, TDM, EG Session 2, R-Squared = 0.331

| | Estimate | Std. Error | df | t value | Pr(>\|t\|) |
|---|---|---|---|---|---|
| (Intercept) | 0.36 | 0.18 | 2.90 | 2.05 | 0.14 |
| condS | 1.27 | 0.09 | 31.14 | 13.60 | 0.00 |
| groupEG2 | 0.61 | 0.43 | 145.08 | 1.42 | 0.16 |

Table 9: Full model, Fixed effects coefficients, language network, FND, EG Session 1, R-Squared = 0.1265

| | Estimate | Std. Error | df | t value | Pr(>\|t\|) |
|---|---|---|---|---|---|
| (Intercept) | 0.44 | 0.12 | 13.59 | 3.57 | 0.00 |
| condS | 0.33 | 0.12 | 31.52 | 2.81 | 0.01 |
| groupEG1 | 0.02 | 0.58 | 176.10 | 0.03 | 0.98 |
| condS:groupEG1 | -0.77 | 0.35 | 13836.93 | -2.22 | 0.03 |

Table 10: Ablated model, Fixed effects coefficients, language network, FND, EG Session 1, R-Squared = 0.1262

| | Estimate | Std. Error | df | t value | Pr(>\|t\|) |
|---|---|---|---|---|---|
| (Intercept) | 0.44 | 0.12 | 13.59 | 3.60 | 0.00 |
| condS | 0.32 | 0.12 | 31.50 | 2.76 | 0.01 |
| groupEG1 | -0.37 | 0.55 | 145.32 | -0.67 | 0.50 |

Table 11: Full model, Fixed effects coefficients, language network, FND, EG Session 2, R-Squared = 0.1271

| | Estimate | Std. Error | df | t value | Pr(>\|t\|) |
|---|---|---|---|---|---|
| (Intercept) | 0.44 | 0.12 | 13.92 | 3.58 | 0.00 |
| condS | 0.33 | 0.12 | 31.53 | 2.80 | 0.01 |
| groupEG2 | -0.56 | 0.58 | 176.50 | -0.96 | 0.34 |
| condS:groupEG2 | -1.05 | 0.35 | 13836.93 | -2.98 | 0.00 |

Table 12: Ablated model, Fixed effects coefficients, language network, FND, EG Session 2, R-Squared = 0.1265

| | Estimate | Std. Error | df | t value | Pr(>\|t\|) |
|---|---|---|---|---|---|
| (Intercept) | 0.44 | 0.12 | 13.91 | 3.61 | 0.00 |
| condS | 0.32 | 0.12 | 31.51 | 2.74 | 0.01 |
| groupEG2 | -1.08 | 0.55 | 145.32 | -1.96 | 0.05 |

Table 13: Full model, Fixed effects coefficients, MD network, FND, EG Session 1, R-Squared = 0.2296

| | Estimate | Std. Error | df | t value | Pr(>\|t\|) |
|---|---|---|---|---|---|
| (Intercept) | 0.71 | 0.23 | 15.54 | 3.03 | 0.01 |
| condH | 1.19 | 0.16 | 31.39 | 7.27 | 0.00 |
| groupEG1 | -0.17 | 0.96 | 57.37 | -0.17 | 0.86 |
| condH:groupEG1 | -0.06 | 0.40 | 6696.94 | -0.14 | 0.89 |

Table 14: Ablated model, Fixed effects coefficients, MD network, FND, EG Session 1, R-Squared = 0.2296

| | Estimate | Std. Error | df | t value | Pr(>\|t\|) |
|---|---|---|---|---|---|
| (Intercept) | 0.71 | 0.23 | 15.53 | 3.03 | 0.01 |
| condH | 1.19 | 0.16 | 31.26 | 7.27 | 0.00 |
| groupEG1 | -0.19 | 0.94 | 52.56 | -0.21 | 0.84 |

Table 15: Full model, Fixed effects coefficients, MD network, FND, EG Session 2, R-Squared = 0.2295

| | Estimate | Std. Error | df | t value | Pr(>\|t\|) |
|---|---|---|---|---|---|
| (Intercept) | 0.71 | 0.23 | 15.99 | 3.05 | 0.01 |
| condH | 1.19 | 0.16 | 31.39 | 7.25 | 0.00 |
| groupEG2 | -0.11 | 0.96 | 57.37 | -0.12 | 0.91 |
| condH:groupEG2 | -0.42 | 0.40 | 6696.94 | -1.05 | 0.30 |

Table 16: Full model, Fixed effects coefficients, MD network, FND, EG Session 2, R-Squared = 0.2295

| | Estimate | Std. Error | df | t value | Pr(>\|t\|) |
|---|---|---|---|---|---|
| (Intercept) | 0.71 | 0.23 | 15.97 | 3.06 | 0.01 |
| condH | 1.19 | 0.16 | 31.26 | 7.21 | 0.00 |
| groupEG2 | -0.32 | 0.94 | 52.56 | -0.34 | 0.73 |

### Supplemental Information III

Values accompanying the Crawford test (a Bayesian assessment of the atypicality of a single-case (EG) score against a set of control scores (Crawford Garthwaite, 2007)). We report statistics for EG's two sessions separately: Session 1 (Table 1) and Session 2 (Table 2).

EG's effect size (*sentences* > *nonwords* for the language network (lang), and *hard* > *easy* for the MD network)) was tested against controls. The number of control participants is reported in the "Control N" column. The "Control $\mu$" and "Control $\sigma$" colums refer to respectively the mean and standard deviation of the controls. The "$CI_{lower}$" and "$CI_{upper}$" colums are the 95% credible interval bounds (the Bayesian interval estimate is called a credible interval rather than a confidence interval), which is the interval estimate of the abnormality of the case's score.

The Crawford tests were implemented using the *psycho* package in R (Makowski, 2018).

Table 1: Crawford statistics for EG session 1

| Network & region | P-val | $EG_{effect\ size}$ | $EG_{percentile}$ | Control $\mu$ | Control $\sigma$ | $CI_{lower}$ | $CI_{upper}$ | Control N |
|---|---|---|---|---|---|---|---|---|
| Lang FDM | 0.237 | 1.855 | 76.358 | 1.361 | 0.687 | 0.182 | 0.293 | 145 |
| Lang TDM | 0.003 | 2.774 | 99.771 | 1.261 | 0.534 | 0 | 0.006 | 145 |
| Lang FND | 0.053 | -0.445 | 5.176 | 0.33 | 0.476 | 0.03 | 0.082 | 145 |
| MD FND | 0.453 | 1.138 | 47.672 | 1.194 | 0.957 | 0.386 | 0.5 | 52 |

Table 2: Crawford statistics for EG session 2

| Network & region | P-val | $EG_{effect\ size}$ | $EG_{percentile}$ | Control $\mu$ | Control $\sigma$ | $CI_{lower}$ | $CI_{upper}$ | Control N |
|---|---|---|---|---|---|---|---|---|
| Lang FDM | 0.201 | 1.941 | 80.065 | 1.361 | 0.687 | 0.15 | 0.256 | 145 |
| Lang TDM | 0.031 | 2.269 | 97.059 | 1.261 | 0.534 | 0.014 | 0.05 | 145 |
| Lang FND | 0.015 | -0.717 | 1.389 | 0.33 | 0.476 | 0.005 | 0.027 | 145 |
| MD FND | 0.335 | 0.779 | 33.225 | 1.194 | 0.957 | 0.236 | 0.44 | 52 |

FDM: Frontal language-dominant; TDM: Temporal language-dominant; FND: Frontal non language-dominant.
